# A Subcortical Model for Auditory Forward Masking with Efferent Control of Cochlear Gain

**DOI:** 10.1101/2024.01.18.576173

**Authors:** Braden N. Maxwell, Afagh Farhadi, Marc A. Brennan, Adam Svec, Laurel H. Carney

## Abstract

Previous physiological and psychophysical studies have explored whether feedback to the cochlea from the efferent system influences forward masking. The present work proposes that the limited growth-of-masking (GOM) observed in auditory-nerve (AN) fibers may have been misunderstood; namely, that this limitation may be due to the influence of anesthesia on the efferent system. Building on the premise that the unanesthetized AN may exhibit GOM similar to more central nuclei, the present computational modeling study demonstrates that feedback from the medial olivocochlear (MOC) efferents may account for GOM observed physiologically in onset-type neurons in both the cochlear nucleus and inferior colliculus (IC). Additionally, the computational model of MOC efferents used here generates a decrease in masking with longer masker-signal delays similar to that observed in IC physiology and in psychophysical studies. An advantage of this explanation over alternative physiological explanations (e.g., that forward masking requires inhibition from the superior paraolivary nucleus) is that this theory can explain forward masking observed in the brainstem, early in the ascending pathway. For explaining psychoacoustic results, one strength of this model is that it can account for the lack of elevation in thresholds observed when masker level is randomly varied from interval-to-interval, a result that is difficult to explain using the conventional temporal-window model of psychophysical forward masking. Future directions for evaluating the efferent mechanism as a contributing mechanism for psychoacoustical results are discussed.

**Significance Statement:** The simulations presented here demonstrate that a recent computational model of the auditory subcortex including medial-olivocochlear efferents generates forward masking, an increase in detection threshold for a short probe tone following a preceding sound. This model explains results from physiological recordings and suggests potential connections to psychoacoustic experiments. The theory that efferent control of cochlear gain is a contributing mechanism for forward masking has several advantages. This theory can explain the strength of masking exhibited by cochlear nucleus neurons, a phenomenon not explained by current physiological theories in which the strength of forward-masking is not increased relative to the periphery until later in the ascending pathway. Additionally, this theory explains results for a psychoacoustic task with random variation in masker level, results not explained by the theory that persistent masker energy interferes with detection of the probe.

## Introduction

Forward masking is the elevation of threshold for detection of a short probe tone due to a preceding sound (masker). This phenomenon has been studied using both physiology (e.g. Relkin and Turner, 1988) and psychoacoustics (e.g. Moore, 1988). At the level of the inferior colliculus (IC), Nelson et al. (2009) demonstrated that physiologically-observed forward masking is often due to a reduction in probe response, rather than interference from a persistent masker response. The present study proposes medial-olivocochlear (MOC) efferent gain control contributes to this reduction.

Neural mechanisms hypothesized to explain decreased probe responses include adaptation of the synapse between the inner hair cell (IHC) and the auditory nerve (AN) (Smith, 1977; Oxenham, 2001) and inhibition of the inferior colliculus (IC) (Salimi et al., 2017; Gai, 2016). The efferent system has also been proposed (physiological recordings: Relkin and Turner, 1988; inference of physiology based on psychophysics: Jennings, 2021; Krull and Strickland, 2008).

However, physiological growth of masking (GOM, or increased probe threshold with increasing masker level) reported for the AN, which should show the effects of the efferent system, is much less than masking observed at later neural stages. Specifically, the range of forward-masked thresholds in anesthetized AN is only 15-20 dB (Relkin and Turner, 1988; Turner et al., 1994), whereas Nelson et al. (2009) and Ingham et al. (2016) report some onset-type IC responses that exhibit GOM closer to 0.5 dB per dB increase in masker level across a masker level range of 80 dB. This result, and neural thresholds with increasing time delay between masker and probe, are roughly similar to results found in psychophysical forward masking tasks (Plack and Oxenham, 1998; Oxenham and Plack, 1997).

Several studies have considered the possibility that probe suppression in the IC arises from the brainstem’s superior paraolivary nucleus (SPON; Nelson et al., 2009; Felix et al., 2015; Gai, 2016; Gao and Berrebi, 2016; Gao et al., 2017; Salimi et al., 2017). However, in at least one experiment (six neurons) chemical inactivation of the SPON did not remove suppression of probe responses for onset neurons, which demonstrate GOM similar to that in psychophysics (Felix et al., 2015, their Fig. 8A). Additionally, Ingham et al. (2016) found GOM in cochlear nucleus (CN) onset neurons comparable to GOM in IC onset neurons, suggesting a potential neural source earlier than the midbrain.

The source of this suppression may be clarified by considering anesthetic effects. IC recordings (Nelson et al., 2009; Ingham et al., 2016) were made in awake and urethane-anesthetized animals; whereas AN recordings (Relkin and Turner, 1988; Turner et al., 1994) used a combination of barbiturate and urethane anesthesia (Dial in urethane). The results of Guitton et al. (2004) demonstrate that efferent activity is decreased more by barbiturate anesthesia than by urethane anesthesia, suggesting that differences in GOM observed in the AN versus IC may be attributable to different anesthetic states rather than different stages of auditory processing (see also Boyev et al., 2002). Greater GOM might be observed in AN fibers if recordings were made using non-barbiturate anesthesia.

Consequently, reconsideration of the efferent system as a contributing mechanism for physiologically-observed forward masking is warranted. We demonstrate that a subcortical model with MOC efferents (Farhadi et al., 2023) explains GOM in neurons that primarily respond to onsets in the IC and CN (Ingham et al., 2016).

We further consider the merits of the efferent mechanism as a contributing mechanism for forward masking observed in psychoacoustics. Psychoacoustic forward masking is often assumed to be caused by masker energy persisting into a temporal window, during which energy from the probe is integrated (Moore, 1988; Oxenham & Plack 1998). Here we consider one specific challenge for this conventional temporal-window model of forward masking: stable thresholds in the presence of a randomly varying masker level (Jesteadt et al., 2005). The present model can account for this result. Further work may evaluate this model on other sets of psychoacoustic forward masking data.

## Materials and Methods

### Stimuli

The stimuli were tone forward maskers followed by short probe tone signals. Stimuli were matched to Ingham et al. (2016), Nelson et al. (2009), or Jesteadt et al. (2005) (Table 1). For Jesteadt et al. (2005), in the 0-σ rove condition, the masker was always 70 dB SPL. In the 6-σ rove condition, the masker level was randomly selected from a Gaussian distribution with a mean of 70 dB SPL and a standard deviation of 6 dB. The distribution was limited to +/- 3 standard deviations. While Ingham et al. (2016) and Nelson et al. (2009) matched the masker and probe frequencies to the best frequency of the neuron, an IC neuron with a characteristic frequency (CF, the frequency to which a neuron is most sensitive) of 4 kHz was modeled here for simplicity. GOM modeling used the stimulus of Ingham et al. (2016) because that study reported GOM data across a range of masker levels for the onset cell-type specifically, whereas Nelson et al. (2009) reported data combined across cell types.

**Table 1:** Table of Stimulus Parameters. Parameters for the stimuli replicated from Ingham et al. (2016), Nelson et al. (2009), and Jesteadt et al. (2005).

| Parameter | Ingham et al. (2016) | Nelson et al. (2009) | Jesteadt et al. (2005) |
| --- | --- | --- | --- |
| Masker Tone Duration | 100 ms | 200 ms | 200 ms |
| Masker Level | Varied re Threshold | 40 dB re Threshold | 70 dB SPL; roved |
| Masker Ramps | 2 ms | 10 ms | 2 ms |
| Probe Tone Duration | 25 ms | 20 ms | 10 ms |
| Probe Ramps | 2 ms | 10 ms | 5 ms |
| Masker-Probe Delay | 2.83 ms | Varied 0-150 ms | 0 and 30 ms |
| Masker Frequency | Varied, 4 kHz modeled | Varied, 4 kHz modeled | 4 kHz in on-frequency condition;<br>2.4 kHz in off-frequency condition |
| Probe Frequency | Varied, 4 kHz modeled | Varied, 4 kHz modeled | 4 kHz |

### Models

The modeling approach was similar to that used for predictions of forward-masked thresholds in Brennan et al. (2023). Two models of the AN were used to simulate neural responses with (Farhadi et al., 2023) and without (Zilany et al., 2014) gain control by MOC efferents. Both AN models had frequency selectivity based on measurements in humans (Ibrahim & Bruce, 2010; Shera et al., 2002) and a CF of 4 kHz. Time-varying AN responses provided the inputs to an IC model cell (Mao et al., 2013) that had a band-enhanced modulation transfer function (MTF, average discharge rate as a function of modulation frequency) with a best modulation frequency of 64 Hz, the median value reported for a population of IC band-enhanced neurons (Kim et al., 2020). Band-enhanced neurons, which were the model IC neurons in results presented here, have average discharge rates that are elevated over a band of modulation frequencies in comparison to the average discharge rate in response to an unmodulated stimulus. Masked tone-detection thresholds were calculated as described below based on the discharge rates of both model AN high-spontaneous-rate (HSR) fibers and the same-frequency inhibition-excitation (SFIE; Nelson and Carney, 2004) model for IC neurons. Note that the IC model of Mao et al. (2013) projected to the MOC, while the final model used for calculating IC thresholds was the SFIE model of Nelson et al. (2004). Mao et al. (2013) is a bandpass modulation filter that provides easily predictable responses to modulation and thus works well as a component of the efferent model. However, the Mao et al. (2013) simple filter model rings after stimulus offset, a property that is not typical in IC neurons. The SFIE model, which does not have this ringing property, was used for the final threshold calculations.

The MOC stage in the model with efferent control of cochlear gain (Farhadi et al., 2023) receives two inputs (Fig. 1): Input 1 is a wide-dynamic range (WDR) feedback signal that controls cochlear gain such that inner hair cells (IHCs) saturate in response to sound levels that are relatively high compared to the spectrum as a whole. This signal was simulated using a low-spontaneous-rate fiber (LSR) AN fiber; as the LSR fiber discharge rate increases, cochlear gain decreases. Input 2 is a fluctuation-driven signal from the IC that decreases cochlear gain for fluctuating channels.

**Figure 1:**
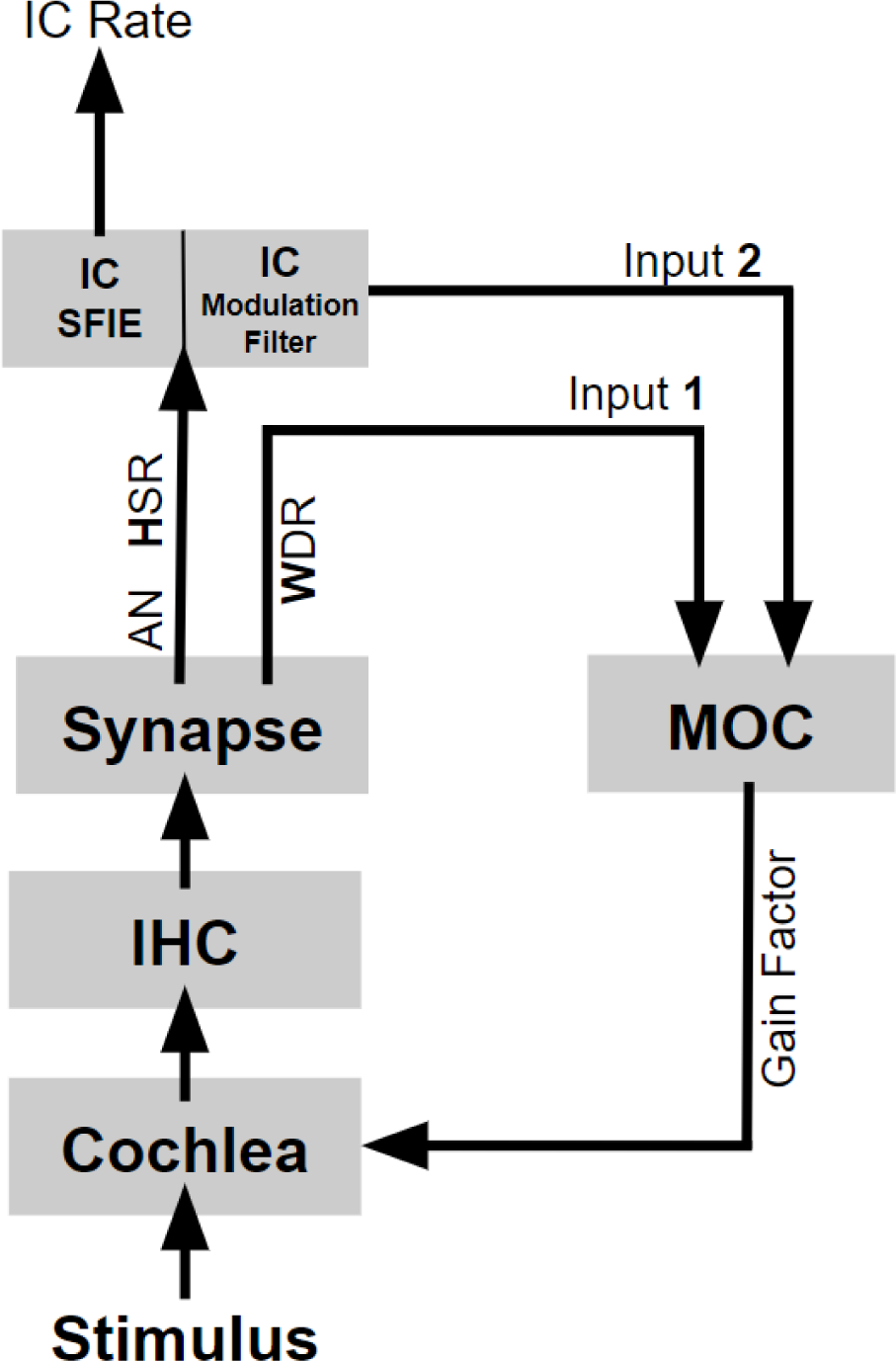
Schematic diagram of subcortical model with MOC Efferents. (adapted from Brennan et al., 2023). Cochlear gain is decreased if the input level (and therefore the WDR discharge rate) is increased. Cochlear gain is also decreased if IC discharge rate is increased by fluctuations in HSR fiber responses. The IC modulation-filter model of Mao et al. (2013) projected to the MOC, while the final model used for calculating IC thresholds was the SFIE model (Nelson et al., 2004).

The process by which the efferent model produced GOM is shown in the blue curves of Fig. 2. Note that the HSR average discharge rate in response to the tone masker was similar during the 40- and 70-dB SPL maskers (left and right columns, respectively), because the average rate of the HSR fibers saturated at or below 40 dB SPL. In contrast, the WDR average discharge rate during the masker was substantially greater at 70-dB SPL (right). In the MOC efferent model of Farhadi et al. (2023), higher WDR rates decrease cochlear gain. Therefore, the gain value decreased more during the 70-dB SPL masker (right) than during the 40-dB SPL masker (left). This difference in gain caused a greater decrease in average discharge rate during the probe when the probe followed the higher-level masker, a pattern that is visible in both the early (HSR and WDR) and more central (IC) responses (Fig. 2). For the model without efferents (gray), the probe response is still present (unmasked) following the 70-dB SPL masker. Note also that the feedback from the IC (“input 2” in Fig. 1) played a relatively small role in these simulations due to the lack of fluctuations in tone maskers.

**Figure 2:**
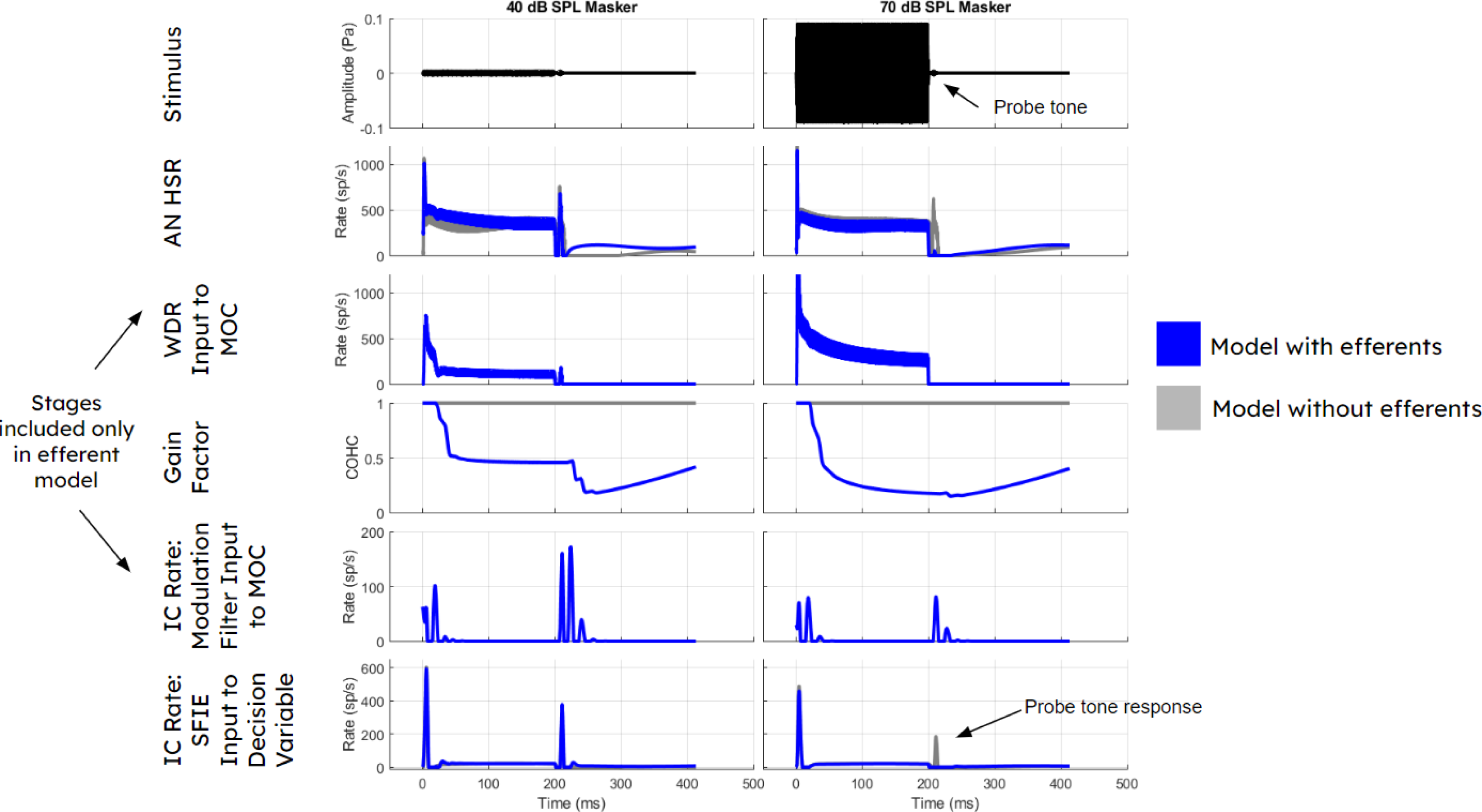
Impact of tone masker level on response to probe tone. The level of a 4-kHz tone masker (left and right columns, 40 and 70 dB SPL) influences the response at various stages of the efferent model (blue). Responses of model without efferents are shown for comparison (gray). Rows show stimulus and five stages of the model: HSR AN fibers, WDR responses (modeled using LSR AN fibers), the gain factor or influence of both feedback pathways on cochlear gain over time; the response of the fluctuation pathway (mostly irrelevant for tone maskers that do not induce fluctuations); and the final response of the IC SFIE model. The LSR stage and feedback pathways are not included in the model without efferents. Note that, for the efferent model (blue), the HSR and IC responses to the probe tone (40 dB SPL, 10 ms, 2.8 ms delay) are substantially decreased for the higher-level masker in comparison to the probe tone responses for the lower-level masker condition, due to a change in cochlear gain. However, for the 70-dB SPL masker, the probe tone responses remain prominent in the model without efferents [gray]).

### Code Accessibility

Matlab code used for the simulations presented here will be made available at https://osf.io/ey4ht/.

### Statistical Methods: Threshold Calculation

Estimated thresholds were based on simulations of a two-interval, forced-choice method-of-constant-stimuli paradigm. Model responses were used to decide, for each trial, whether the tone was in interval 1 or 2. For the physiological simulations, on each trial, the interval with the higher maximum response during a 40-ms time window was selected as the target interval. The use of maximum response during a time window was based on the modeling approach of Brennan et al. (2023) and causes results to depend on the onset response of the model neuron (essentially converting the SFIE model response, which has minimal steady-state response and a strong onset component, into an onset neuron model). The time window used for the physiological simulations matched the analysis windows used in Nelson et al. (2009) and Ingham et al. (2016), and is shown as a short solid horizontal line in Fig. 3. This window started 5 ms after the beginning of the probe tone stimulus so that, taking into account a few milliseconds of response latency from the auditory nerve model, the window included most of the probe response while excluding the masker response. This window placement is in comparable to the windows in Nelson et al. (2009) and Ingham et al. (2016). In most conditions, the masker and probe responses did not overlap in time, matching the lack of persistence of the masker response reported by Nelson et al. (2009).

**Figure 3:**
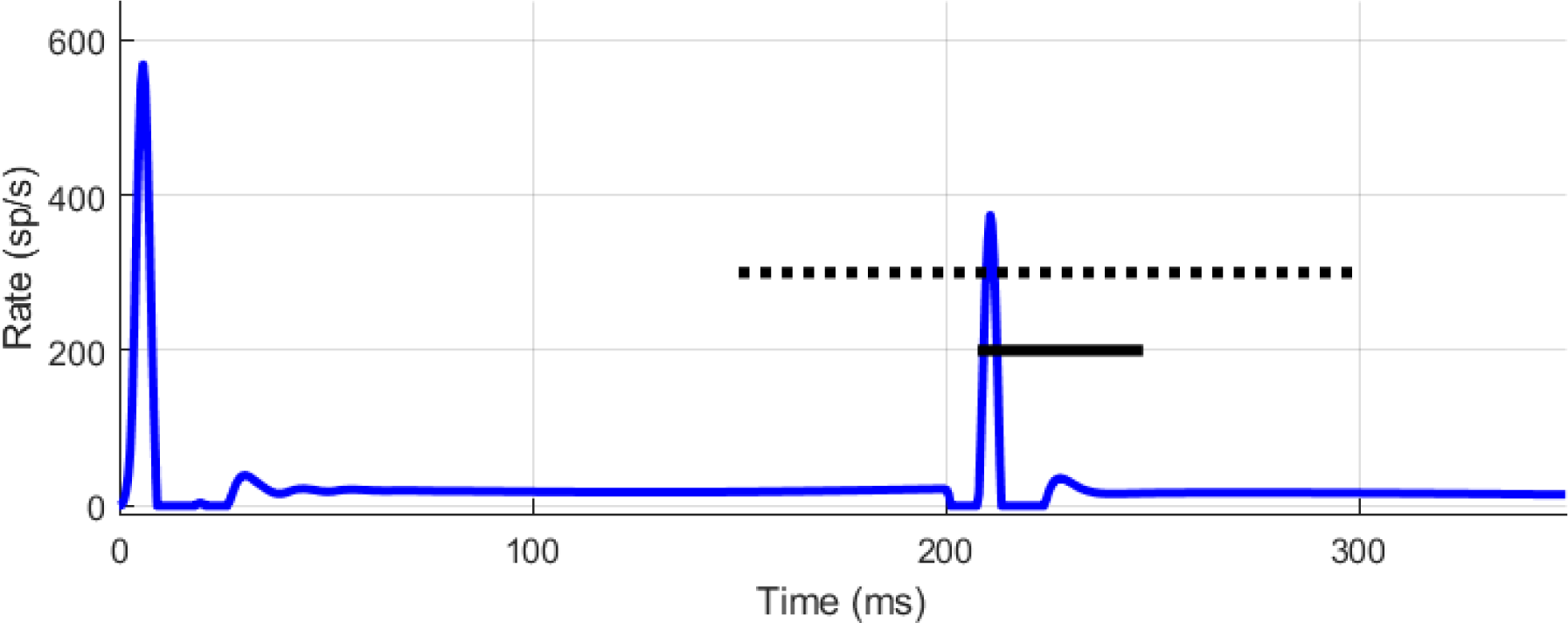
Decision-variable windows. IC SFIE model response to a 4 kHz, 40-dB SPL masker, 2.83 ms masker-probe delay, and 40 dB SPL, 10-ms probe tone (IC received input from model with efferents). Lines superimposed to show the placement of the physiological decision-variable window (bottom solid line) and psychophysical decision-variable window (top dotted line). The physiological decision-variable window was used for simulating the results of Ingham et al. (2016) and Nelson et al. (2009), and was shifted later or earlier depending on the masker-probe delay. The psychophysical decision-variable window was only used for simulating the paradigm of Jesteadt et al. (2005), and was not shifted with the masker-probe delay.

For the psychophysical simulations, this procedure was the same except that the decision-variable window extended from 150 to 300 ms to reflect less a priori knowledge about the time of the probe response (Fig. 3: long dotted horizontal line). This window was not shifted in time for the different delays of 0 and 30 ms.

Percent correct was tabulated over 100 trials per target level. A logistic curve was fitted to the data across tone levels to estimate threshold (61% to match Ingham et al. (2016), 70.7% in all other cases). Note that the time windows used here differed from windows used in the temporal-window model of forward masking (Moore et al., 1988). Here, the windows were time spans over which response maxima were detected, rather than windows over which energy was integrated.

### Specifying Thresholds in SPL vs. SL

Unit conversion was necessary for comparing the simulation results to physiological data because masker levels in the physiological papers (Ingham et al., 2016; Nelson et al., 2009) were described relative to neuron threshold (SL; suppression due to masker), whereas the model requires input to be scaled to a specific SPL. The following four points describe our approach: (1) For modeling the physiological data, we treated the unmasked threshold of the model neuron as analogous to the unmasked threshold of the neuron recorded in the experiment, and matched simulated masker levels relative to the unmasked threshold of the model neuron (SL). (2) For modeling the psychoacoustic task with randomly varying level, masker levels in the simulation were matched to the SPLs used in the original experiment (Jesteadt et al., 2005). (3) It was possible to compare model thresholds to both psychoacoustic and physiological data because the model neuron unmasked probe tone thresholds were close to 0 dB SPL. Therefore, SPL and SL were roughly equivalent for the model. The greatest deviation from an unmasked threshold of 0 dB SPL was approximately 5 dB SPL for the AN model (with and without efferents) with the probe tone parameters used by Nelson et al. (2009). The IC model (with and without efferents) in the same condition had an unmasked threshold close to 1 dB SPL. (4) We looked for psychoacoustic data to compare to the model performance in the physiological tasks, to better understand whether one mechanism could explain both types of data. We chose data from a task that was as similar as possible to the physiological tasks (Moore and Glasberg, 1983). The masker-level conditions chosen for comparison to the model data were matched to the SPL used in the simulation. This choice allowed for realistic application of level-dependent model features, such as frequency selectivity and saturation of the synapse between the IHC and AN.

Although the points above describe the matching of masker level (simulation input) in the simulations, a separate question is whether to compare thresholds (simulation output) to physiological and psychoacoustic thresholds in terms of SPL or SL. It is reasonable to assume that some additional noise would elevate both unmasked and masked thresholds for the system as a whole, compared to a single-neuron model with the optimal CF for detecting the signal. Therefore, thresholds are expressed in SL (suppression due to masker) in all cases. In cases where psychoacoustic thresholds were expressed in SPL in the original paper without reporting unmasked thresholds, thresholds were converted from SPL to SL by subtracting 15 dB. This conversion was used by Oxenham and Plack (1998) for stimuli similar to those used here.

## Results

### Growth of Masking

Growth of masking in AN and IC Models with efferents (Fig. 4; blue squares, blue diamonds, respectively) was similar to physiological recordings of IC onset-type neurons from Ingham et al. (2016; black solid line). The IC model with efferents inherited suppression properties from the AN model with efferents, as demonstrated by the similarity between IC and AN model results with efferents (Fig. 4: blue symbols). GOM was substantially greater for the models with efferents (blue) than the models without efferents (gray). The IC models generally had higher thresholds than the AN models. GOM was similar between (1) the models with efferents (blue symbols), (2) physiological recordings of onset-type neurons from Ingham et al. (2016; black solid line), and (3) psychophysical results reported by Moore and Glasberg (1983; dashed line) for similar probe tone duration, masker-probe delay, and SPLs similar to those used for the models here (note that the psychophysical data is matched to the model simulations based on SPL rather than SL; see Methods). Also note that CN onset-type neurons in Ingham et al. (2016) showed similar GOM to IC neurons, and thus a fit to the IC data implies a reasonable fit to the CN data, as the efferent GOM mechanism described here would have effects before the CN in the ascending pathway. However, the types of CN neurons observed in Ingham (2016) were not explicitly modeled here.

**Figure 4:**
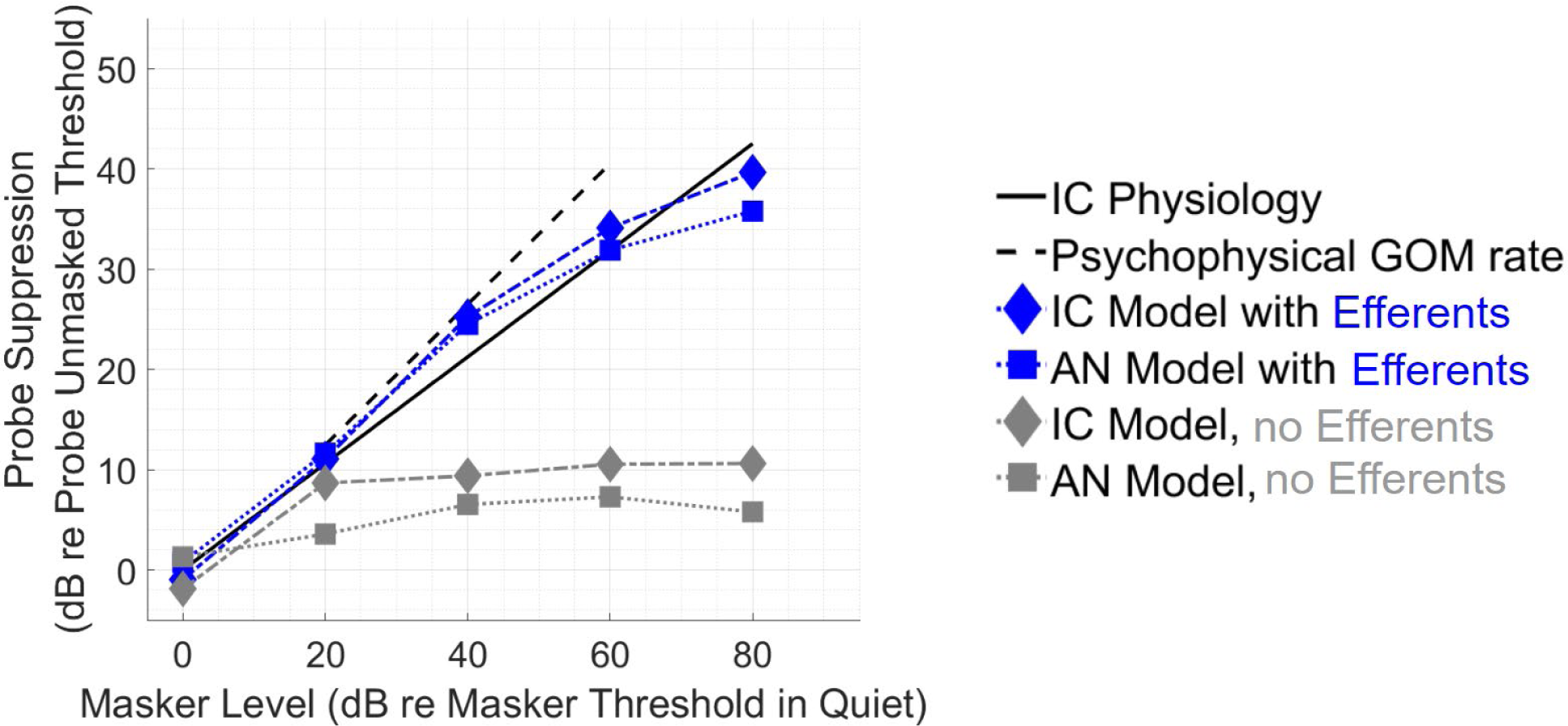
Forward-masked thresholds vs. masker level for models, physiology, and psychophysics. Growth of masking thresholds shown for IC and AN models (diamond and square markers, respectively), with and without efferents (blue and gray, respectively; color online). IC Physiology (black solid line) based on line fit to average suppression in onset-type neurons from Ingham et al. (2016): 2-ms delay, 100-ms masker, 25-ms signal. Psychophysical GOM rate (slope of black dashed line) derived from Moore and Glasberg (1983): 0-ms delay, 210-ms masker, 20-ms signal. For physiology and models, masker level is plotted on the x axis relative to the threshold of the masker in quiet. Probe suppression is plotted on the y axis, in dB relative to the unmasked threshold of the probe. Psychophysical GOM curve is shifted along the x axis to more closely match the SPL of model stimuli (SL and SPL were similar for the model because model unmasked thresholds were within 5 dB of 0 dB SPL). The IC physiological results shown here (from Ingham et al, 2016) were from experiments using urethane anesthesia, with potentially less impact on the efferent system than barbiturates (Guitton et al., 2004).

### Change in Masked Threshold with Masker-Probe Delay

In the physiological data of Nelson et al. (2009), masked threshold decreased by 19 dB for masker-signal delays between 0 and 150 ms (Fig. 5; black solid line). Without efferents, the AN model threshold decreased by 6 dB (gray squares), and the IC model threshold decreased by 11 dB (gray diamonds). Efferents brought this decrease in masked thresholds closer to the physiological range, leading to a decrease of 15 dB for the AN model (blue squares) and 19 dB for the IC model (blue diamonds). Suppression in the IC model with efferents was close to that of IC physiological recordings at short and long delays, but did not decrease as a function of delay rapidly enough at moderate delays. The IC model with efferents inherited most suppression properties from the AN model with efferents, although there were slight differences between the AN and IC. Relative to the AN, the IC sharpened the response to the probe at short delays because of sensitivity to modulation, and therefore the IC had lower thresholds than the AN in SPL (not shown; Fig. 5 shows suppression relative to unmasked threshold).

**Figure 5:**
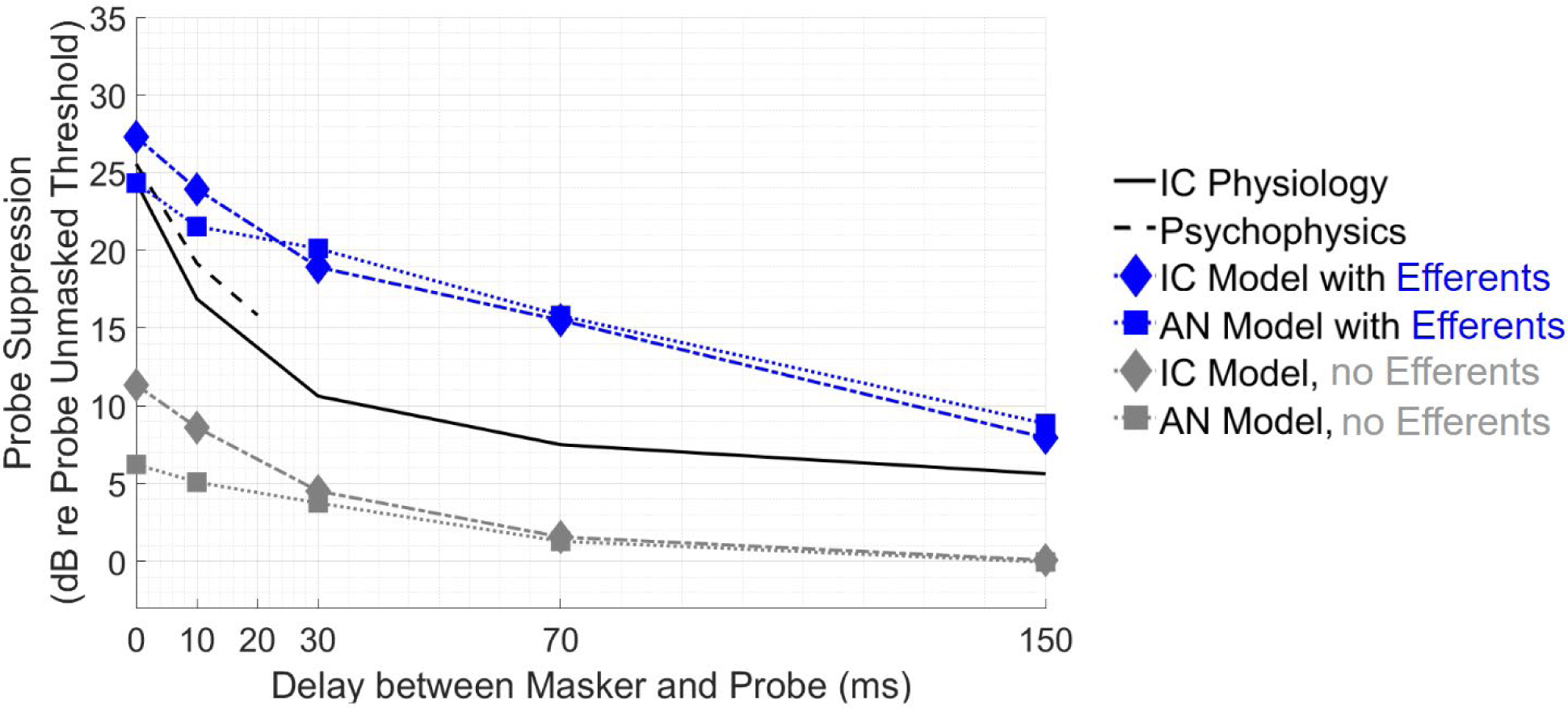
Change in Threshold with Increasing Masker/Probe Delay. Suppression at varying delays shown for IC and AN models (diamond and square markers, respectively), with and without efferents (blue and gray, respectively). IC Physiology (black solid line): average suppression in awake recordings of IC neurons in Nelson et al. (2009): masker was 40 dB re neuron threshold (SL), 200-ms in duration, 20-ms signal. Psychophysics: from Moore and Glasberg (1983): masker at 40-dB SPL, 210-ms masker, 20-ms signal. Simulations matched parameters in Nelson et al. (2009); unmasked model thresholds were about 1 dB SPL for the IC models and about 5 dB SPL for the AN models, so 40-dB SPL and 40 dB SL were similar values for the models.

This IC/AN difference applied for both models with and without efferents. However, the unmasked threshold of the AN was also about 4 dB higher than the IC (unmasked thresholds: IC with efferent: 1 dB SPL; IC without efferent: 0.8 dB SPL; AN with efferent: 5 dB SPL; AN without efferent: 5 dB SPL). This difference in unmasked thresholds caused suppression to be lower in the AN and obscured the enhanced encoding in SPL provided by the IC. Psychophysical and physiological data (dashed and solid black lines, respectively) were similar at short delays, and the IC model with efferents showed a similar decrease in threshold over these delays.

### Masked threshold with roving-level masker

The previous results demonstrated the capacity of the Farhadi et al. (2024) IC model with efferents to account for aspects of physiological masking data, which in some cases is similar to psychoacoustic masking data. We also challenged the model with a psychophysical result from Jesteadt et al. (2005) that is difficult for the temporal window model (Moore et al., 1988; Oxenham and Plack, 1998) to explain.

Jesteadt et al. (2005) reported that randomly varying (roving) the level of the masker between intervals did not increase thresholds, regardless of whether the masker was on-frequency (masker and probe at 4 kHz) or off-frequency (masker at 2.4 kHz, probe at 4 kHz). The on-frequency result is consistent with the temporal window model if cochlear nonlinearities are taken into account, as cochlear compression or AN discharge rate saturation should cause the neural response to the masker to change little across the range of randomly varied levels when the masker is at the same frequency as the probe, and this consistent masker response should prevent the masker-level rove from impacting the detection task. However, the off-frequency result is contrary to the prediction of the temporal-window model, as the neural response to the masker should change with the roving level: according to that model, higher masker energy in the standard interval should cause listeners to mistake that interval for the target (a false alarm), and lower masker energy in the target interval should cause listeners to mistake that interval for the standard (a miss), thus increasing listener thresholds when the masker level is roved. Jesteadt et al. (2005) also observed that listener decisions in a roving-level forward-masking task were correlated with the masker level in the target interval but not the masker level in the standard interval, which again suggests that the false alarms and misses described above were not occurring in the way that the temporal-window model would predict. Although other models featuring persistence of masker energy, such as temporal template models (the temporal pattern of masker + probe response, stored in memory), can explain this result, the conventional temporal window model cannot (Jesteadt et al., 2005). Here we tested whether the present model could explain these results without a temporal template stored in memory.

For the following tests, the model decision process considered the response over a long duration, from halfway through the masker until the end of the stimulus interval. This window length differed from the short time windows used for modeling the physiological data, which were short to match the duration of time windows used to record spikes in the physiological studies (see Fig. 3). The window length was extended for the simulations of human psychoacoustics based on the assumption that human listeners may not be able to precisely attend to activity within the short time windows used to measure discharge rate in the physiological studies.

Thresholds for the on- and off-frequency maskers based on the IC model with efferents did not change when a roving masker level was introduced (pairs of blue open vs. blue filled symbols), matching this difficult-to-explain feature of the psychophysical data (pairs of black open vs. black filled symbols). Even in the off-frequency masker condition, for which the temporal window model would predict a substantial rove effect, the IC model with efferents showed minimal effect of the rove, similar to the psychophysical data. For the on-frequency masker result, model thresholds were close to psychophysical thresholds in the on-frequency, 0-ms delay, and in the off-frequency, 30-ms delay conditions, but were elevated compared to psychophysical thresholds for the on-frequency, 30-ms delay condition, similar to Fig. 5. These thresholds reflect too much suppression in the model at that delay. The model also provides too little suppression at the 0-ms delay for the off-frequency masker (right panel).

This lack of rove effect (matching the psychophysical data) occurred in both the on-frequency and off-frequency conditions because the IC model had minimal steady-state response to the tone masker (shown in Fig. 2), a property consistent with its use here to represent onset-type neurons. The false alarms and misses that would occur due to the roving masker level in the temporal-window model did not occur in this model because a higher masker level in the standard was not reflected in the steady-state response of the neuron, so a higher masker level would not be mistaken for a higher probe response. Likewise, the standard interval never caused a miss due to having a higher steady-state masker response than the target. Of course, the masker level in the target interval still influenced probe detection by producing changes in gain. Higher masker levels decreased gain, decreased the probe response, and decreased percent correct in some target intervals; lower masker levels increased gain, increased the probe response, and increased percent correct in some other target intervals. These cases balanced out (just as they would if the psychophysical task only involved one interval, instead of two) and yielded the same threshold as without rove, because the ratio of hits to false alarms was the same. Considering all of these factors together, model performance in this task depended only on the target interval, and not on the standard interval. This result aligns with the finding of Jesteadt et al. (2005) that the masker level in the target interval was correlated with whether trials were correct or incorrect, whereas the masker level in the standard interval was not. As mentioned above, Jesteadt et al. (2005) noted that their results were explainable if listeners matched a temporal template to the shape of the masker-probe response in the target interval. A temporal template and the IC decision process used here are similar in that they are robust to the roving-level paradigm. However, the present neural implementation has the advantage of not requiring any memory template.

It should be noted that thresholds for AN HSR model fibers (not shown) were *also* stable in the roving-level paradigm; however, this stability occurred for more complex reasons. In the case where masker and probe were both at 4 kHz, thresholds remained the same in the roving-level paradigm because the AN HSR fibers were saturated and changes in masker level were not reflected in the masker response. This result was consistent with the expected outcome hypothesized by Jesteadt et al. (2005). In the off-frequency condition, in which the masker was at 2.4 kHz, the response to the masker steady-state did change with level; however, the change in the steady-state response to the masker between 70 and 88 dB SPL was small compared to the growth in the probe response near threshold (therefore, minimal false alarms occurred for trials with high masker levels in the standard interval), and the decrease in the steady-state response to the masker below 70 dB SPL did not create misses because the decision variable detected maximum discharge rate (and therefore always recorded the probe response, and not response to the masker, in those conditions). AN HSR responses to the off-frequency condition would not have been robust to the roving masker level if the simulated decision variable had been based on mean rates instead of maximum rate; however, the IC response would have still been robust to the roving level with either decision variable, because the IC model onset response contributed much more to the IC average discharge rate than did the IC steady-state response. Thus, although the masking itself was provided by the efferent feedback, the success of the IC model in simulating robustness to roving masker level was largely due to the IC model’s lack of a steady-state response.

## Discussion

### Summary

The results above demonstrated that simulations based on a recent subcortical model (Farhadi et al., 2023) can explain multiple aspects of physiological forward-masking data. The MOC efferent feedback in this model added substantial GOM in the AN responses to a forward-masking paradigm, compared to a prior AN model (Zilany et al., 2014). This GOM was inherited by the IC model. The IC model had GOM similar to that in physiological recordings of onset-type cells in the IC (Nelson et al., 2009; Ingham et al., 2016) and of onset-type cells in the CN (Ingham et al., 2016), with some similarities to human psychophysical results (Moore and Glasberg, 1983). This model had more forward masking across masker-probe delay than the Zilany et al. (2014) model, which was based on AN responses under anesthesia. The model presented here also displayed a timecourse of changes in forward-masking thresholds that was similar to that in IC physiology (Nelson et al., 2009). Finally, the lack of response to the steady-state of the masker in IC onset cells explained the stability of forward-masked thresholds in a paradigm with a randomly varying masker-level, a result that is difficult to explain using the temporal-window model of forward masking (Jesteadt et al., 2005).

### Do the MOC efferents provide a plausible underlying mechanism for physiologically-observed forward masking?

Previous work suggested that physiological forward-masking varies depending on the response type of the neuron (e.g., onset, sustained, etc.) (Ingham et al., 2016). Here we focused on the responses of onset-type neurons in the IC, partially motivated by similarities between IC onset-type responses to forward masking and human psychophysics. Physiological studies outline constraints for the underlying masking mechanism: (1) the mechanism may influence forward-masking at both CN and IC stages (Nelson et al., 2009; Ingham et al., 2016); (2) there is no evidence that the mechanism is reflected in the response of the barbiturate-anesthetized AN, perhaps due to suppression of the efferent system (Guitton et al., 2004, Relkin and Turner 1988; Turner et al., 1994), although the amount of forward masking present in the AN when the MOC efferent system is active is unknown; (3) the mechanism must decrease the response to the probe rather than temporally extending the response to the masker; that is, persistence of the masker response is rarely observed at the level of the IC (Nelson et al., 2009). Of course, the masker response may still persist, or overlap with the response to the probe, in later processing stages whose responses are shaped by slower time constants.

The mechanism of efferent-driven changes in gain fits all of these constraints: The efferent system is capable of influencing forward masking at both CN and IC stages because it influences cochlear gain; this system would be deactivated in the barbiturate-anesthetized AN, explaining the lack of physiological GOM observed by Relkin and Turner (1988; also Turner et al., 2004); and this system would decrease the response to the probe rather than temporally extending the response to the masker. Note that forward masking observed in onset-cell types specifically requires probe suppression rather than persistence (regardless of whether psychophysical decisions are influenced by persistence), as these neurons have a minimal sustained response.

However, the proposed mechanism of MOC efferent-driven gain control raises important questions for future investigation: why, in the studies by Nelson et al. (2009) and Ingham et al. (2016), was there greater growth of masking for onset-cell types than other cell types, when efferent-driven cochlear gain control could hypothetically influence all cell types? Are the transient onset responses affected differently by efferents than steady-state responses? Additionally, the model has too much suppression at moderate delays between 10 and 70 ms (Fig. 5), an aspect that is reflected in the excess suppression at 30 ms in Fig. 6. This and other inaccuracies may be fixed in future model iterations by taking into account the combined effect of other sources of suppression (inhibition from SPON or synaptic adaptation at the CN or IC level).

**Figure 6:**
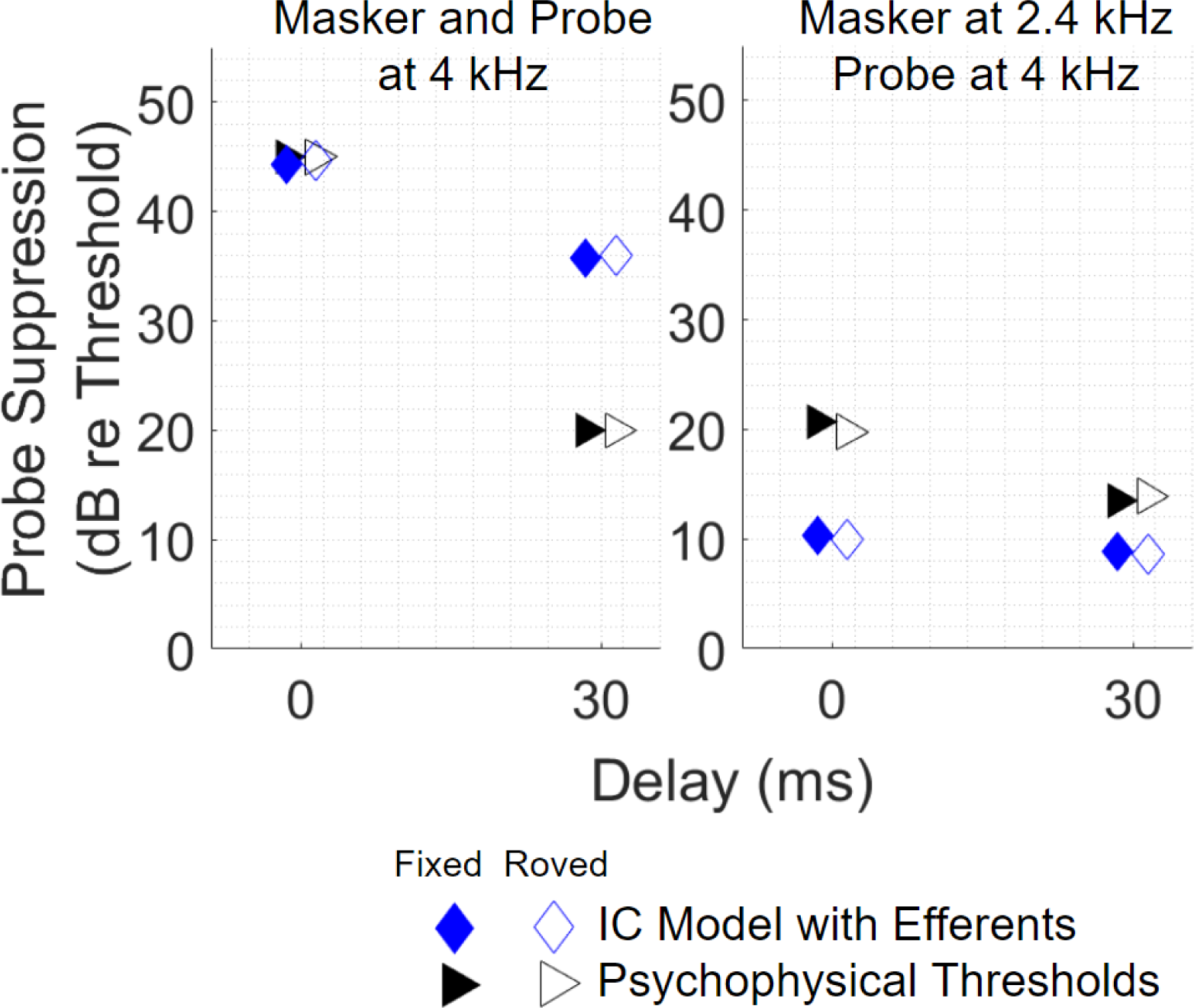
Stable Thresholds with Masker-level Rove. Psychophysical thresholds (black triangles; Jesteadt et al., 2005: experiment 1) and thresholds for IC model with efferent feedback (blue diamonds) shown for masker-probe delays of 0 and 30 ms, with masker level fixed at 70 dB SPL and with masker-level rove (filled and open symbols, respectively). Left panel: on-frequency condition (masker and probe at 4 kHz). Right panel: off-frequency condition (masker at 2.4 kHz, probe at 4 kHz). Pairs of points with similar thresholds indicate the minimal impact of rove (x-coordinate offset to clearly show both filled and open symbols). Psychophysical suppression was calculated by subtracting a general estimate of unmasked thresholds for similar probe tones (15 dB SPL; Oxenham and Plack, 1998) from data reported in Jesteadt et al. (2005). Simulations matched parameters in Jesteadt et al. (2005).

### Is it plausible that physiological forward masking, and specifically the MOC-efferent mechanism, can explain forward masking observed in psychoacoustic studies?

As an explanation for psychophysics, the efferent-gain mechanism has at least two strengths. First, as discussed above, the mechanism is suitable for explaining physiological results that bear similarities to psychophysical results. Figures 4 and 5 show several nontrivial similarities between physiological and psychophysical forward masking, including similar growth of masking. That said, these comparisons include only a small sample of psychophysical forward masking data, and the model produces too much suppression at moderate delays as discussed above.

Second, this model can explain the stability of thresholds with roving masker level in the results of Jesteadt et al. (2005), which pose a problem for the temporal window model of forward masking (Moore et al., 1988; Oxenham and Plack, 1998; Jesteadt et al., 2005). If the temporal-window model were correct, when the masker level was randomly varied, the persistent energy at the time of the probe should also vary, making the energy contributed by the probe more difficult to detect and increasing the thresholds. However, thresholds remain unchanged in this condition. The level of the masker must still be encoded, but variations in level do not interfere with detection of the probe. The use of onset-type cells to encode the presence of the probe tone offers a solution to this problem. With minimal representation of the steady-state response of the masker, it does not matter whether the level of the masker is roved up or down in the standard interval; the masker level does not cause false alarms (though properties of the masker can still affect masking if they change the activity of the efferents). Without interference from the standard interval, the cases of higher and lower gain in the target interval balance out, leaving thresholds unchanged in the roving condition. This mechanism also explains why correct/incorrect answers by participants in Jesteadt et al. (2005) were correlated with masker energy in the target interval, but not with masker energy in the standard interval.

One question is why the cognitive algorithm for detecting the probe tone should use the output of onset neurons (which have greater GOM) to detect the probe, when there are other types of neurons with less GOM and therefore, potentially more salient probe responses at high masker levels. On the other hand, this disadvantage is outweighed by the fact that onset neurons do not encode the sustained response to the masker, and thus onset-type responses are pre-processed to focus on the relevant details for an algorithm that detects the beginning of a second sound (the probe). The Jesteadt et al. (2005) experiment is just one demonstration of how useful it is to ignore irrelevant aspects of the masker for the purpose of probe-tone detection.

While this theory offers a solution to the finding of Jesteadt et al. (2005), whether it can explain psychoacoustic forward masking in general remains to be seen. Some circumstances where forward masking occurs pose challenges for this model. For example, forward masking exists with similar properties for listeners with cochlear implants (Shannon, 1990), where the efferents cannot play a role. Similar properties also occur for backward masking (Oxenham and Moore, 1994) and for masker-probe combinations that are only 25 ms in length, most likely too brief for the efferents to take effect (Yasin et al., 2013). A different kind of challenge is presented by the finding that masking decreases for longer probe tones (Oxenham, 2001), which is difficult to explain if the relevant cells only encode the probe-tone onset, a property tied to substantial GOM in physiology and robustness to level rove in our simulations. On the other hand, it is possible that some onset neurons with substantial GOM may also include very limited sustained response (Ingham et al., 2016). Interestingly, forward masking due to phenomena that emerge binaurally such as Huggins pitch (Plack et al., 2013), may be possible to account for using this theory, as in this view the efferents receive feedback from IC neurons with bilateral inputs.

It is possible that one or more of these tasks involve a different source of masking from the others or are accomplished using a different decision process from the others: i.e. forward masking in listeners with cochlear implants may involve masking that originates due to the mechanism of electrical stimulation itself, or the roving-level task may involve a different decision process than in more classic tasks. However, there are similarities across these cases that suggest a common mechanism; for example, the time course of forward masking in cochlear implants is similar to that in acoustic hearing (Shannon, 1990). Further psychoacoustic experiments may help address the potential of different mechanisms: for example, if listener performance does not improve with increasing tone duration when the masker level is roved, or if listener performance is no longer robust to rove in that case, that would suggest that the listener’s decision process differed between different forward masking tasks.

Another possibility is that detection by onset-type neurons underlies the lack of influence of masker-level rove in Jesteadt et al. (2005) as proposed here, and efferents have some psychophysical effects (consider separate lines of evidence in Brennan et al., 2023 and Jennings, 2021), but that psychophysical GOM in forward masking is not caused by MOC efferents, accumulating instead from contributions of other neural sources. For example, synaptic adaptation throughout the CN, SO, and IC, which is not included in the present model, may elevate masked thresholds. Evidence already reviewed in this paper suggests that some forward masking exists in the AN even when the MOC efferent system is suppressed, and that the SPON is capable of contributing to forward masking, although neither of these sources can fully account for growth of masking observed in the CN onset-type neurons. In this view, forward masking in cochlear implants (as well as forward masking for short maskers and probes) and GOM in backward masking would be easier to explain: the source of masking would not be removed in cochlear implants or short maskers and probes, and an onset detection mechanism similar to the one proposed here, calculating a max over a given window, could produce backward masking that would increase with the level of the masker (each probe onset would need to compete with a growing masker onset). This view would not attribute to efferents the primary role in masking, but would predict no change in thresholds due to masker-level rove in the experiment of Jesteadt et al. (2005).

In summary, any theory aiming to reconcile physiological and psychoacoustic observations of forward masking must deal with several challenges. The large body of experimental evidence relating to forward masking presents unique resources, but also unique challenges, to model accounts of these phenomena.

## Conclusions

Overall, the simulations presented here support the hypothesis that MOC efferents contribute to forward masking observed physiologically (decreased probe responses observed in the IC) and suggest that efferents may play a role in psychophysical forward masking, though open questions remain. These simulations explain why listener thresholds are robust to randomly varying masker level, a surprising result that the temporal-window model (Moore et al., 1988) cannot predict. Additionally, this surprising result is explained without requiring templates stored in memory, instead using the simple mechanism of onset-type neurons that do not respond to the steady-state of tone maskers. These findings extend previous inquiries into the role of MOC efferents in psychophysical masking (Jennings and Strickland, 2012, 2010; Jennings et al., 2011; Jennings et al., 2009; review: Jennings, 2021). The theory proposed here accounts for the GOM observed in recordings of onset-type neurons in the IC. Furthermore, this theory provides a conceptual account for how GOM could occur in the CN while not occurring in AN recordings performed under barbiturate anesthesia: the efferent system may be less active under that type of anesthesia compared to other anesthetics (Guitton et al., 2004). In summary, this theory provides potential answers to open questions about forward masking stemming from previous experiments in both psychoacoustics and physiology.

## Acknowledgements

Supported by NIH-NIDCD-R01-010813 (BNM, AF, LHC) and a grant from the Hearing Industry Research Consortium (RFP2022) (AS, MB). Braden Maxwell is currently affiliated with the Department of Psychology at the University of Minnesota. Afagh Farhadi is currently affiliated with the Department of Speech, Language, and Hearing Sciences at Purdue University.

